# Modelling the gut microbiota of children with malnutrition: *in vitro* models reveal differences in fermentability of widely consumed carbohydrates

**DOI:** 10.1101/2024.05.08.593150

**Authors:** Jennifer Ahn-Jarvis, Kendall Corbin, Suzanne Harris, Perla Rey-Troncoso, Peter Olupot-Olupot, Nuala Calder, Kevin Walsh, Kathryn Maitland, Gary Frost, Frederick J. Warren

## Abstract

There is increasing evidence in children suffering from Severe Acute Malnutrition (SAM) that there is disruption of the gut microbiome and low gut microbiota diversity, which may be contributing factors to poor outcomes during nutritional treatment and recovery. The gut microbiome of children with SAM has been demonstrated to have a lower production of beneficial short chain fatty acids, which may contribute to impaired gut barrier function. Recently, several microbiota-directed therapies have been tested in clinical trials in children with SAM. Among them we hypothesized that feeds containing fermentable carbohydrates from various sources (legumes, chicory, milk oligosaccharides) would be fermented to produce beneficial microbial metabolites by the microbiota of children with SAM. In this study we used an *in vitro* model system inoculated with stool from children with SAM to investigate the fermentability of four substrates; inulin (a chicory-derived fructan), two milk powders (one supplemented with a human milk oligosaccharide) and a chickpea enriched feed. We demonstrated that while the milk powders and chickpea feed were fermented to produce short chain fatty acids, inulin was only fermented to a very limited degree. Through 16S rRNA sequencing we demonstrated that the samples inoculated with inulin had low microbial diversity and linked this to the limited ability to metabolise inulin. Through revealing the fermentability of different complementary feeds, the findings of this study will be of use for the design of future therapeutic feeds for treatment of SAM.

**Importance:** Malnutrition is a major contributor to childhood mortality globally and is a major public health problem primarily affecting Lower- and Middle-Income Countries. Despite the development of nutritional recovery therapies, for those with the severe and complicated form of malnutrition (SAM), mortality and relapse rates remain high. Emerging evidence suggests a role for the gut microbiome in these poor outcomes, which is known to be significantly altered in children in SAM, compared to healthy age matched controls. To aid in recovery from SAM, nutritional interventions should be designed to support the gut microbiome, using a range of ingredients targeted for colonic fermentation. It is important to understand the fermentation capacity of the gut microbiome of children with SAM, to design future nutritional interventions. In this work, we demonstrate that inulin, a widely used chicory-derived prebiotic, is not a suitable fermentation substrate for the gut microbiome of SAM children, while legume-based formulations and milk oligosaccharides result in increased production of beneficial metabolites.

## Introduction

Undernutrition is still a widely prevalent problem in many Lower- and Middle-Income Countries (LIMCs), affecting almost 25% children under the age of 5 years globally, and being implicated in almost half of all childhood deaths (1, 2). Childhood undernutrition can be broadly split into two categories, namely stunting (length-for-age Z-score [LAZ] ≤2) and wasting (weight-for-height Z-score ≤2) (1). Although less prevalent than stunting, the most severe form of wasting, Severe Acute Malnutrition (SAM) is associated with higher mortality rates. For children hospitalised with SAM up to 20% die, with a high proportion of deaths occurring during the first week of hospitalisation (3). Nutritional rehabilitation alongside supportive treatments has been the cornerstone of SAM treatment. The feeding regimes have been developed to deliver optimum nutrition to children hospitalized with SAM. The current WHO recommendation for nutritional treatment of acutely sick children hospitalized with SAM are mainly milk-based feeds (called F75 and F100 indicating their calorie content/100ml) for inpatient management followed by Ready to Use Therapeutic Feeds (RUTF) as a paste for rehabilitation. This treatment regime leads to weight gain and recovery of appetite, as well as improvement in other indicators such as glycaemic control (4). Despite this, nutritional recovery (as well as recovery of anthropomorphic indicators) in SAM is a poor indicator of long term outcomes (5, 6), indicating that the damage caused by SAM is far more complex and long-lasting than simple nutritional deficiencies.

The human gut microbiota has recently become an area of intense research interest, driven by advances in metagenomic sequencing technologies (7, 8), but also a greater understanding that the gut microbiota influences many physiological processes including; nutrient acquisition (9); growth hormone signalling and appetite control (mediated through production of Short Chain Fatty Acids (SCFA)) (10); and immune regulation (11). During long term recovery from SAM there is evidence of systematic dysregulation of many of these processes, leading to altered appetite regulation impaired acquisition of key nutrients and increased susceptibility to a range of infections (12). This points to the potential for a key role for the microbiota in the pathogenesis and rehabilitation of SAM.

16S based amplicon sequencing studies of the stool microbiota of infants suffering from SAM have revealed that there are dramatic alterations in the gut microbiota (13–17). Twins studies on infants discordant for SAM have demonstrated that distinct microbiota changes occur as a result of SAM, independent of host genetics, environment and background diet (13). These changes include a general reduction in α and β diversity, as well as increases in potentially pathogenic bacteria such as *Klebsiella* and Enterobacteriaceae, and a reduction in beneficial saccharolytic bacteria in *Clostridium* clusters IV and XIVa (such as *Blautia*, Lachnospiraceae, Ruminoccoaceae and *Faecalibacterium prausnitzii*), *Bacteroides* species and *Lactobacillus* (15, 18). It has been hypothesised that some aspects of the microbiome of children with SAM represent a failure of the gut microbiota to fully mature (15, 16), although there is also strong evidence that there is significant overgrowth of potential enteropathogens (17, 18), and risk of invasive bacterial infection (3).

Emerging evidence indicates that reversing the gut microbiota changes that occur as a result of SAM have the potential to improve outcomes (19), although gut microbiome targeted therapies have to be carefully designed as interventions using probiotics and synbiotics have had limited success (18, 20). The most promising approach is the use of so called ‘microbiota directed foods’ (21, 22), in which food ingredients rich in carbohydrates that are fermentable by the gut microbiota of children with SAM are supplemented conventional RUTFs. The selection of fermentable carbohydrate is crucial as the damage to the gut microbiota diversity that occurs during SAM has the potential to restrict the range of substrates which are accessible to the microbiota. By successfully identifying substrates that support the metabolism of the broadest range of microbial species possible, an increase in SCFAs may be achieved.

In this study, we aimed to directly determine the fermentability of potential microbiota directed foods by the gut microbiota sampled from children with SAM by employing and *in vitro* batch fermentation model seeded with stool samples collected from infants hospitalised with SAM. We investigated the fermentation of 4 different substrates. Infant formula, similar in composition to the F75/F100 formula was selected as a baseline for the standard SAM recovery formula. Inulin, a chicory-root derive fructan oligosaccharide, was selected as it is a widely used prebiotic carbohydrate (23) which is highly fermentable by the gut microbiota of healthy infants (24), but conversely in animal models of the SAM gut microbiota it was shown to not to support growth (25). The same animal study demonstrated that milk oligosaccharides similar to those found in human milk did support weight gain (25), so we selected a milk powder supplemented with 2’- Fucosyllactose to reflect this. Finally, we used a legume enriched feed designed to support microbial recovery in SAM (26, 27) formulated with chickpea and high in resistant starch (28, 29). Chickpea has been successfully used as a feed supplement to support microbiota recovery in moderate acute malnutrition (19), and the formulation used here has been specially formulated for use in intervention trials to support recovery of infants with SAM (26). Each of these foods was used as a substrate for *in vitro* batch fermentation in models seeded with stool samples from children hospitalized with SAM. Metabolomic analysis of the fermentation media using ^1^H NMR was conducted to indicate substrate fermentability through production of microbial metabolites during the time course of fermentation. Samples were also taken for 16S metataxonomic analysis to determine the effects of each of the substrates on microbial diversity and potential pathogen burden.

## Results

### Cohort demographics and baseline microbiome composition

The demographics of the cohort are shown in Table 1. The average age of the cohort is 2.4 years, with an age range of 0.9 years to 4.5 years. All children included in this study met the criteria for SAM with an average MUAC of 11.1 cm. Clinical data from this cohort was collected from February to April 2016 from day of admission to day 7 of hospitalization.

**Table 1.**
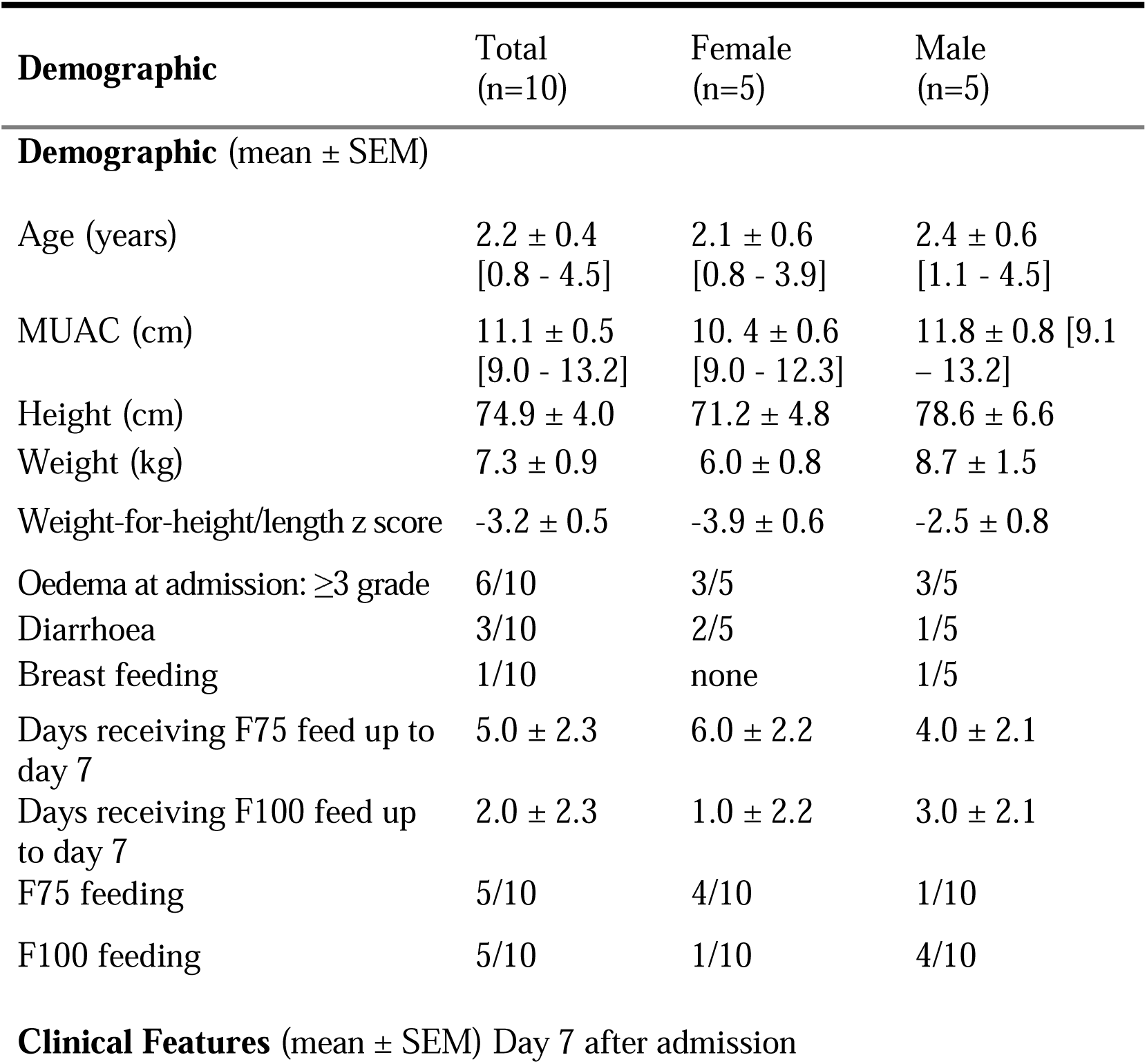
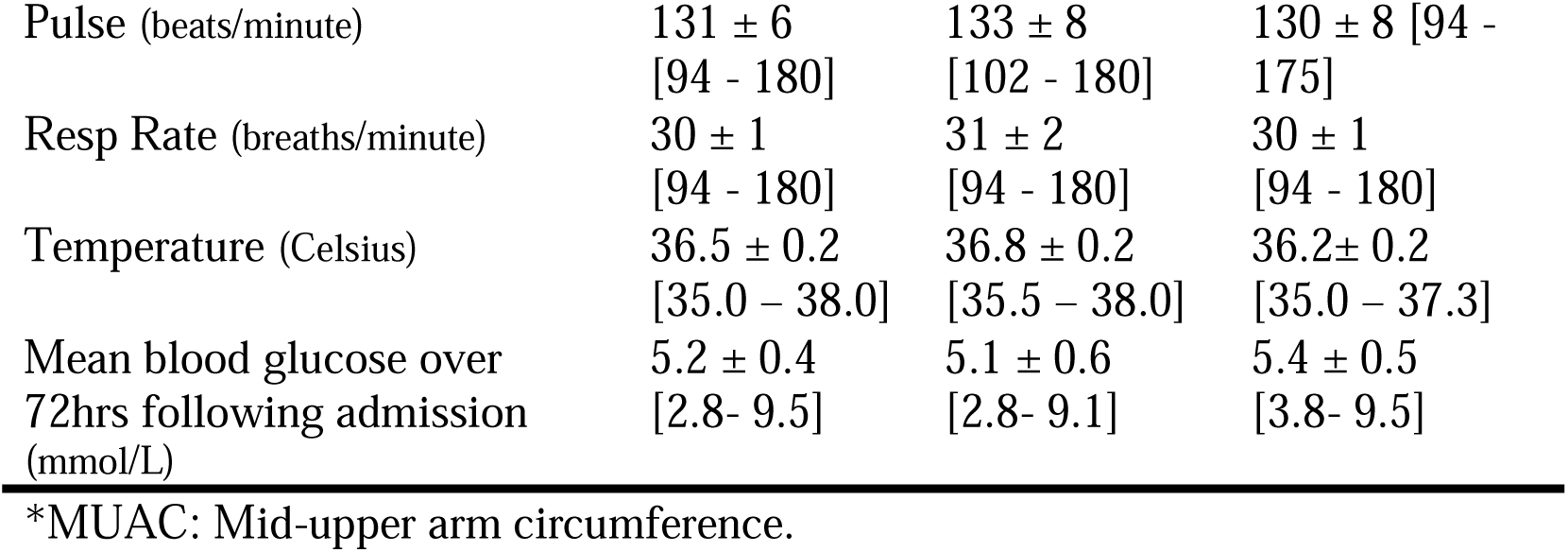
Demographics of the study cohort. Values are given as mean ± standard deviation.

| Demographic | Total<br>(n=10) | Female<br>(n=5) | Male<br>(n=5) |
| --- | --- | --- | --- |
| **Demographic (mean ± SEM)** |  |  |  |
| Age (years) | 2.2 ± 0.4<br>[0.8 - 4.5] | 2.1 ± 0.6<br>[0.8 - 3.9] | 2.4 ± 0.6<br>[1.1 - 4.5] |
| MUAC (cm) | 11.1 ± 0.5<br>[9.0 - 13.2] | 10.4 ± 0.6<br>[9.0 - 12.3] | 11.8 ± 0.8 [9.1<br>– 13.2] |
| Height (cm) | 74.9 ± 4.0 | 71.2 ± 4.8 | 78.6 ± 6.6 |
| Weight (kg) | 7.3 ± 0.9 | 6.0 ± 0.8 | 8.7 ± 1.5 |
| Weight-for-height/length z score | -3.2 ± 0.5 | -3.9 ± 0.6 | -2.5 ± 0.8 |
| Oedema at admission: $\geq 3$ grade | 6/10 | 3/5 | 3/5 |
| Diarrhoea | 3/10 | 2/5 | 1/5 |
| Breast feeding | 1/10 | none | 1/5 |
| Days receiving F75 feed up to day 7 | 5.0 ± 2.3 | 6.0 ± 2.2 | 4.0 ± 2.1 |
| Days receiving F100 feed up to day 7 | 2.0 ± 2.3 | 1.0 ± 2.2 | 3.0 ± 2.1 |
| F75 feeding | 5/10 | 4/10 | 1/10 |
| F100 feeding | 5/10 | 1/10 | 4/10 |
| **Clinical Features (mean ± SEM) Day 7 after admission** |  |  |  |
| Pulse (beats/minute) | 131 ± 6<br>[94 - 180] | 133 ± 8<br>[102 - 180] | 130 ± 8 [94 -<br>175] |
| Resp Rate (breaths/minute) | 30 ± 1<br>[94 - 180] | 31 ± 2<br>[94 - 180] | 30 ± 1<br>[94 - 180] |
| Temperature (Celsius) | 36.5 ± 0.2<br>[35.0 – 38.0] | 36.8 ± 0.2<br>[35.5 – 38.0] | 36.2 ± 0.2<br>[35.0 – 37.3] |
| Mean blood glucose over<br>72hrs following admission<br>(mmol/L) | 5.2 ± 0.4<br>[2.8- 9.5] | 5.1 ± 0.6<br>[2.8- 9.1] | 5.4 ± 0.5<br>[3.8- 9.5] |

\*MUAC: Mid-upper arm circumference.

### In vitro fermentation reveals substrate driven differences in SCFA production

*In vitro* fermentation experiments were established using a batch model colon fermentation system seeded with stool samples from each of the 10 SAM trial participants who were on standard nutritional and supportive treatments. Each of the fermentation vessels included either inulin, MIMBLE feed, SMA or Similac milk powders as substrates. Samples were taken at defined timepoints (with baseline samples, time = 0, taken prior to fermentation), and microbial metabolites were quantified using ^1^H-NMR (Fig. 2, Table S1). Following up to 36 hours of fermentation, all the substrates tested gave rise to production of SCFA’s (p value = 0.007) and other microbial metabolites, with significant differences identified between substrates.

**Figure 1.**
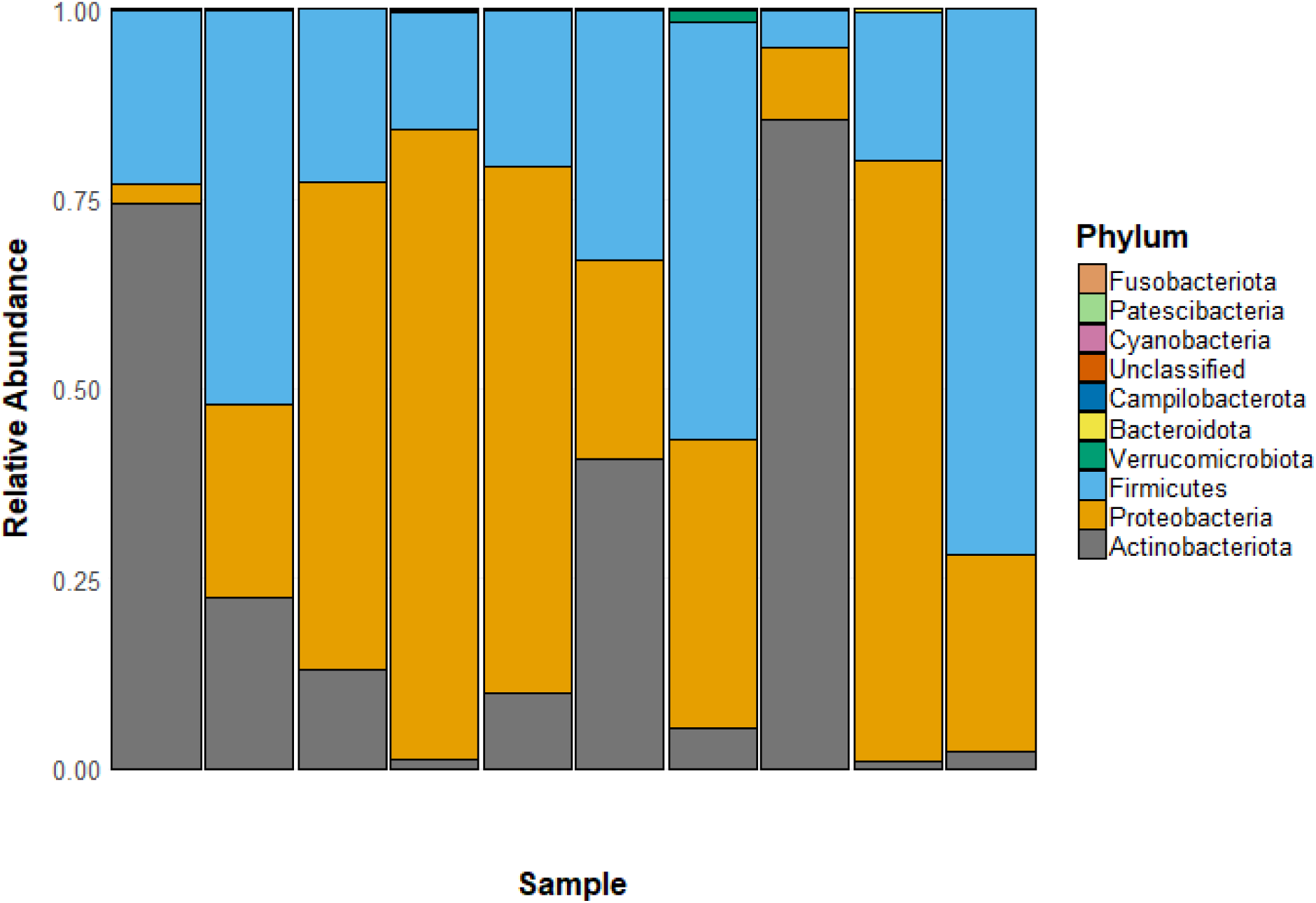
Participant-level phylogeny profile showing the Phylum level abundances of bacterial taxa determined by 16S sequencing at the start of the experiment (time = 0), prior to *in vitro* fermentation.

**Figure 2.**
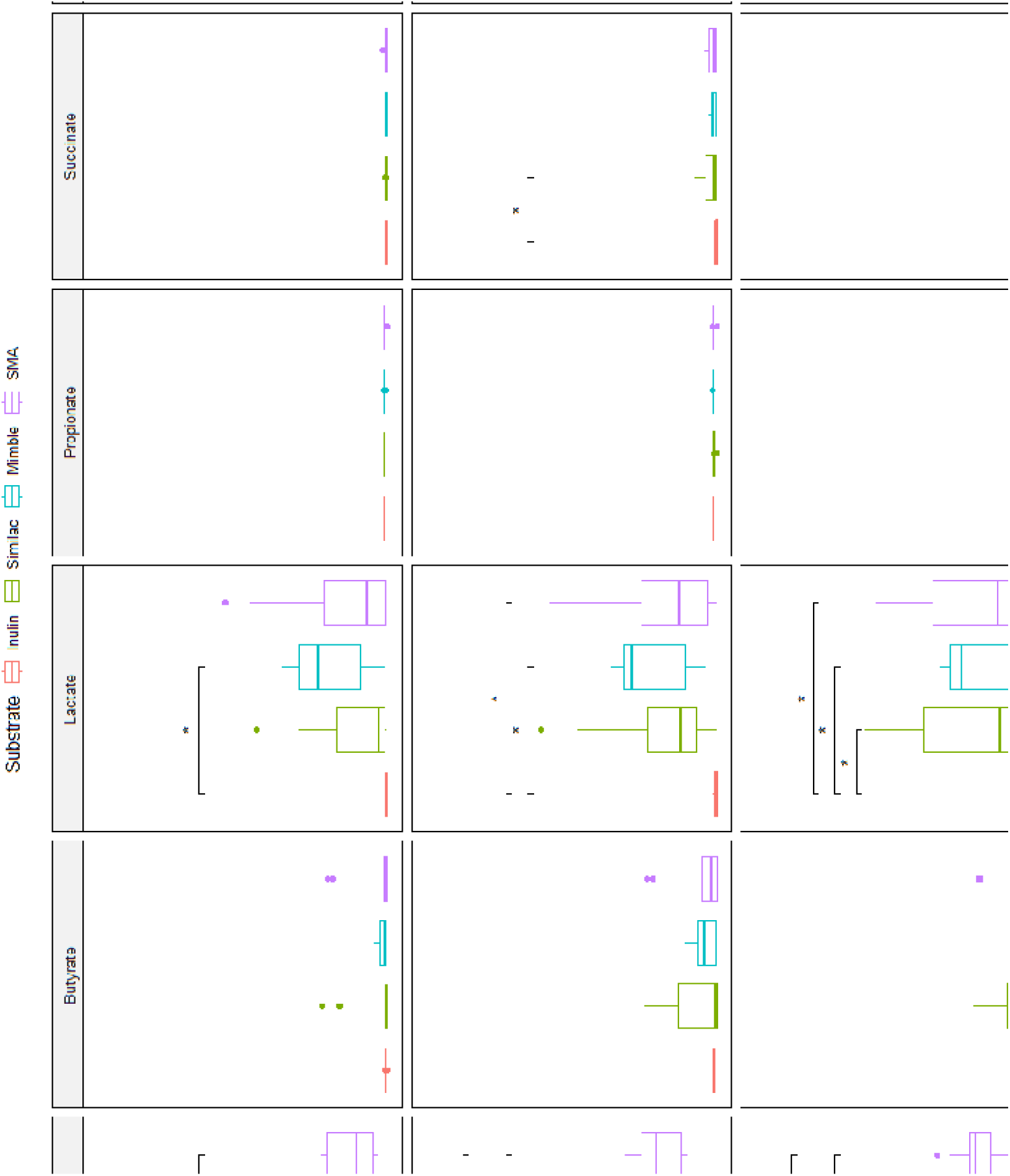
Short chain fatty acid concentrations determined by ^1^H NMR following 12, 24 and 36 hours of *in vitro* fermentation. Metabolite differences were tested using ANOVA with a post-hoc Tukey’s test. Statistically significant differences are indicated with * p-value < 0.05, ** p-value < 0.01, *** p-value < 0.001, **** p-value < 0.0001. All SCFA concentrations are indicated in mM.

The total SCFA production from the inulin substrate was significantly lower than from the other three substrates after 24h of fermentation (Fig. 2). The differences in total SCFA were mainly driven by acetate and butyrate production. Acetate was highest for the two milk powders, slightly lower for the MIMBLE feed and significantly lower for the inulin. Butyrate levels were similar between the MIMBLE, SMA and Similac substrates, but lower in the vessels with inulin.

Propionate production was not significantly different across all substrates and timepoints, although it should be noted that propionate production was low in all the fermentations, and only modest increases from baseline were seen with the MIMBLE and SMA substrates. The clearest differences observed between the substrates were in the metabolic intermediates lactate and succinate. Lactate was produced in response to MIMBLE, Similac and SMA substrates from 12 h of fermentation onwards, while succinate showed delayed kinetics of formation, with the peak of metabolite production occurring between 12 and 24 h (Fig. 2, Table S1). Neither lactate nor succinate were produced in significant quantities from the fermentation of inulin. The absence of these metabolic intermediates would hinder the production of SCFA end products and may be reflected in the lower butyrate and acetate concentrations observed for inulin compared to the other substrates tested in this study (9, 31). It is likely that the low production of microbial metabolic intermediates reflects limited breakdown of inulin by the microbiota inoculated into these models. Evidence for limited breakdown of inulin can be derived from the ^1^H NMR spectra of the fermentation media following 36h of fermentation (**Fig 3**). Several peaks are observed only in the inulin supplemented media which can be assigned to inulin, including the peaks at 4.3 and 4.1ppm which arise from protons in the fructose ring of inulin (32). These inulin peaks remain invariant at all the timepoints, suggesting that the inoculum used in these fermentations is unable to degrade and utilize inulin as a substrate and it remains unmetabolized throughout the course of the fermentation experiments, unlike the three other substrates tested which were fermented at similar rates.

**Figure 3.**
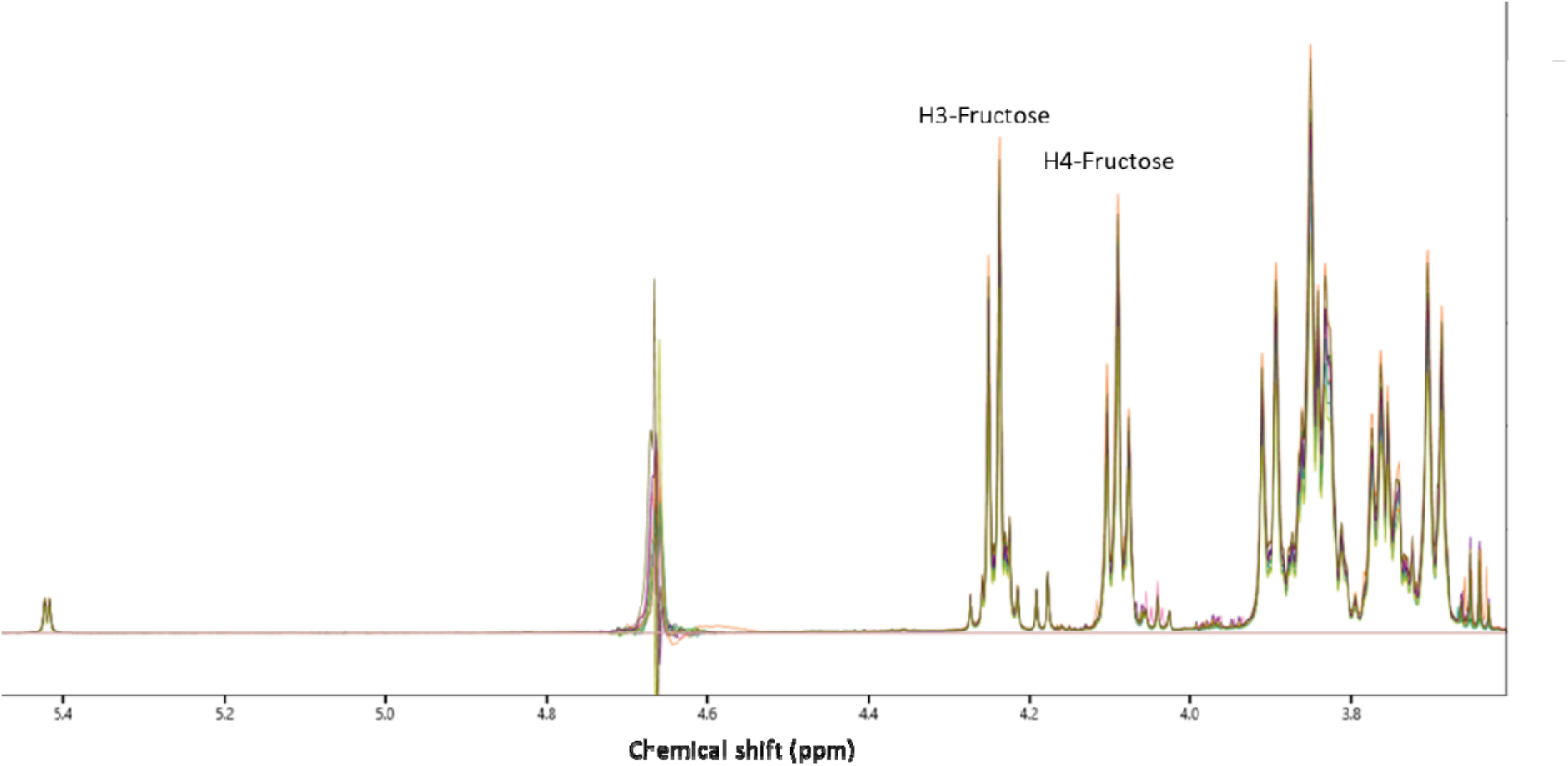
^1^H NMR spectra for fermentation media sampled from each of the inulin substrate fermentation vessels following 36h of fermentation. Peak assignments from Caleffi et al.(32) for the H3 and H4 protons in the fructose ring.

### Substrates drive differences in microbiome composition

The microbial composition within the fermentation vessels (Fig. 4) changed over the time course of the experiment, depending on both time and substrate. Immediately following inoculation, the microbial community composition for each of the substrates was near identical (Fig. 4), with the 0 h timepoints, sampled immediately following inoculation, clustering very closely together on a PCoA plot (Fig. 4) reflecting the individual microbial composition of each donor. Following sampling at subsequent time points the microbiome compositions deviated from the composition at 0h, depending both on the substrate and the time point and by 48 h the microbial community composition has moved away from that seen at baseline, although there were not significant differences observed between the substrates.

**Figure 4.**
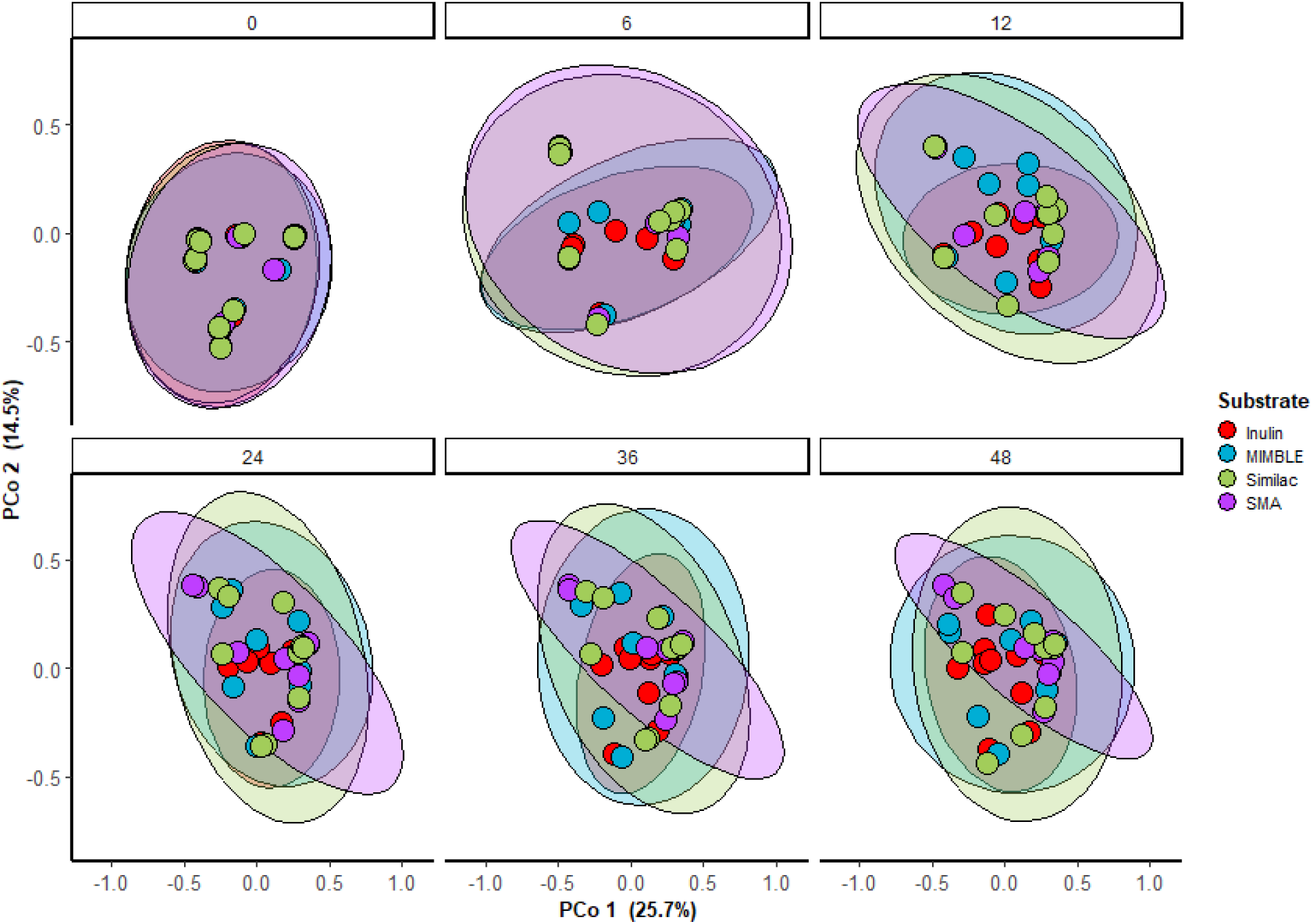
PCoA plot based on Bray-Curtis dissimilarity at ASV level for microbial communities coloured by substrate at 0, 6, 12, 24, 36 and 48 hours of fermentation.

Across all the substrates there was reduction in microbial diversity (Shannon diversity) observed over time in the models, although this did recover slightly at later time points (Fig. 5a). The greatest reduction in microbial diversity was observed in the Similac and SMA milk powders at the 12h time point. In contrast, the Shannon diversity in vessels inoculated with inulin was less reduced at 12h, and at subsequent time points, compared to the other substrates. The changes in microbial diversity are reflected in the phylogeny profiles (**Fig 5b**), where the inulin remains similar to baseline after 24 h fermentation, whereas the other three substrates see reductions in Actinobacteriota and Proteobacteria, and increased Firmicutes abundance.

**Figure 5.**
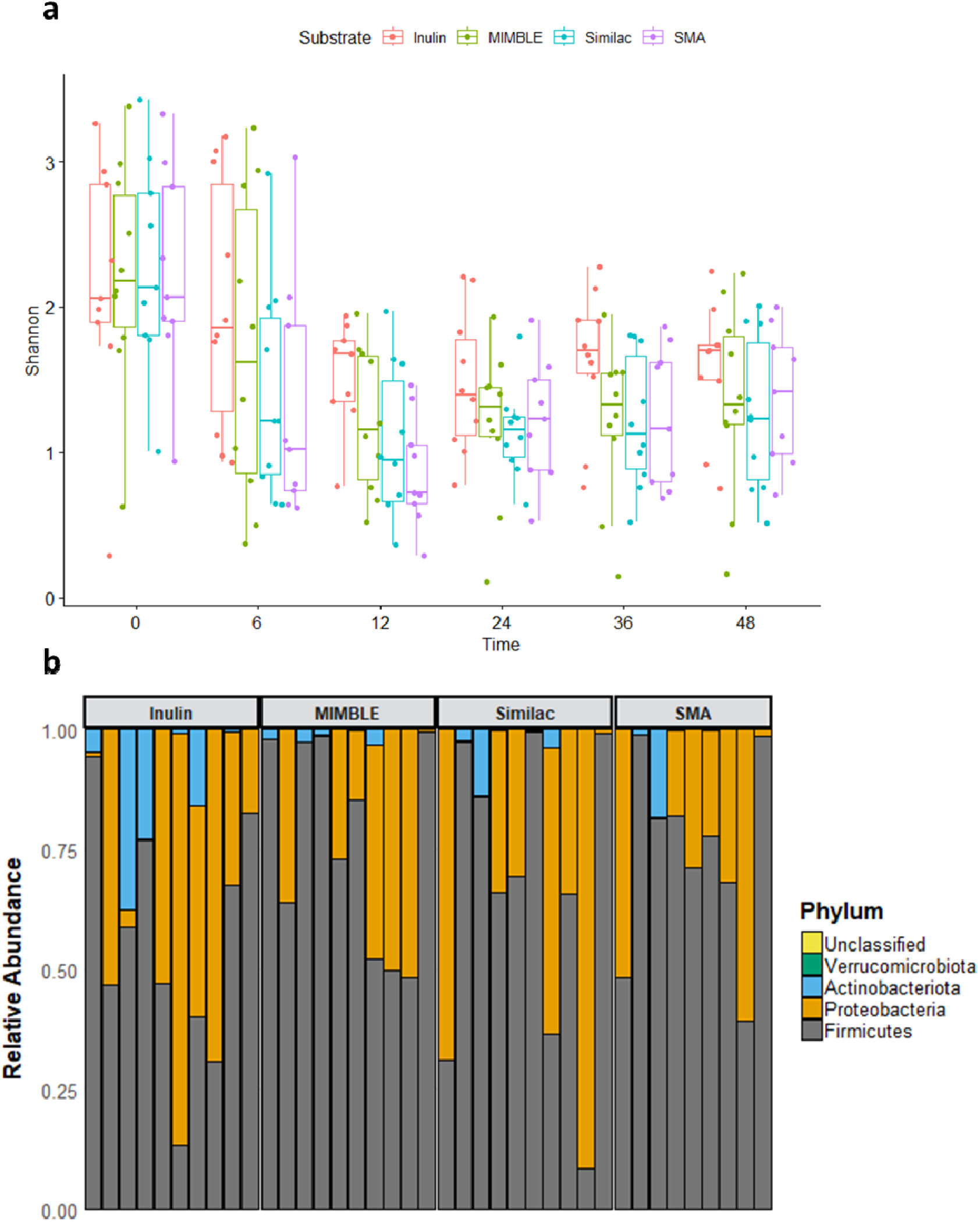
Microbiome changes during fermentation. a. Changes in alpha-diversity (Shannon index) over time for each of the substrates during fermentation. b. Phylogeny plot showing phylum level taxa abundances for each participant fermenting each individual substrate following 24h of fermentation.

## Discussion

The gut microbiome has recently emerged as an important target in the treatment of SAM, due to a series of studies indicating that SAM is associated with gut microbiome immaturity (15, 16, 33), leading to reduced production of beneficial microbial metabolites such as SCFA which are protective of the gut endothelium and are crucial to healthy gut function. In a clinical setting, the gut microbiome can be further damaged by the extensive use of antibiotics in treatment regimes. In several studies, reduced stool SCFA has been associated with increased mortality in children with SAM (27, 34). As a result, dietary interventions building on the standard WHO recommendations to include ingredients which specifically target the gut microbiome, such as chicory inulin and legume based ingredients have been developed and trialled in interventions with SAM patients (21, 26, 27, 35, 36).

In this paper we have adapted an *in vitro* model colon protocol to mimic the gut microbiome of SAM patients. The basal media used is a minimal media with only key vitamins and minerals for microbial growth, but no additional carbon sources reflecting the restricted diet of SAM patients. To establish the gut microbiota of the SAM patients in vitro, stool samples collected from the patients are seeded into the vessels. The stool samples were selected from patients who had been in clinical treatment for SAM for 7 days following a standard WHO treatment pathway, and therefore reflected the impact of SAM, standard WHO milk-based nutritional feeds and antibiotic treatment on the gut microbiota of SAM patients (27). This was reflected in the microbial composition of the samples which were inoculated into the model, which showed high levels of Proteobacteria and very low levels of Bacteroidetes. The high abundance of Proteobacteria was also observed in the larger cohort that these samples were obtained from (27), where Proteobacteria levels were found to peak at 7 days following hospital admission, the point at which the samples in our study were taken. Low relative abundances of Bacteroidetes (<5% in all samples) were observed in these samples, consistent with observations in previous studies of infant malnutrition (15, 37).

Using this model, we demonstrate that inulin is almost completely unfermentable by the gut microbiota sampled from infants with SAM. In previous studies investigating *in vitro* fermentation of inulin by stool sampled from infants, there was a clear age dependent effect where the gut microbiome of very young infants is unable to ferment inulin, while older infants can partially ferment lower molecular weight inulin (24, 38). These studies, however, investigated much younger infants than the current study (median age 2.2 years). This may reflect the immaturity of the gut microbiome of children with SAM. In adults, short chain inulin’s are fermented preferentially by species of the genus *Bifidobacteria* while longer chain inulin’s are fermented preferentially by *Bacteroides* (39). The low levels of Bacteroidetes in the samples analysed in this study, and in children with SAM, may limit the fermentability of inulin in these groups. Recent animal model studies and an intervention study in children with SAM found similar results, with inulin failing to increase faecal SCFA or increase weight gain (25, 27). The mechanism underpinning this is not clear from the data presented in this study.

*Bifidobacteria* were observed to be more abundant in the fermentation vessels containing inulin compared to the other substrates, although the difference was not statistically significant (Fig. S1). However, *Bacteroides* species were not observed to increase in abundance in the inulin fermentation vessels (Fig. S2). The reasons for the lack of growth of *Bacteroides* species are not known. In contrast, the ‘MIMBLE’ feed, which is enriched with legumes, and milk powders both with and without human milk oligosaccharides, were found to be fermentable. Animal studies have demonstrated that the human milk oligosaccharide (HMO) 2’-FL contributes to increased SCFA output and supported weight gain in a model of SAM (25). In the present study the fermentability of the two milk formulas tested was very similar, and the SCFA output of both was high. In contrast to HMO’s, legumes are cheap and widely produced, therefore making an excellent candidate for addition to supplementary feeds for use in the treatment of SAM. In the present study we tested the MIMBLE feed, containing 10% chickpea flour, which was found to be fermented by the gut microbiota of children with SAM in an *in vitro* model, yielding a range of SCFAs. The chickpea enriched feed also resulted in a greater reduction in Proteobacteria abundance during fermentation than the other substrates tested. Several recent studies have identified that feeds containing chickpea flour have the potential to act as a food source for the gut microbiota of children with SAM, due to the diverse range of polysaccharides including pectins and arabinoxylans present in the cell wall polysaccharides of chickpea. These polysaccharides can be used to modulate the gut microbiome and promote the growth of beneficial bacteria in the immature gut microbiota of SAM patients (19, 35, 36). These results indicate the potential for the use of locally sourced plant-based foods, such as chickpeas, for use in recovery foods which can support SCFA production in the colon as effectively as HMO’s (21, 35, 40).

In conclusion, in this study we have used an *in vitro* model of the gut microbiome of children with SAM to test the fermentability of four different substrates. Sequencing revealed that the microbiome of the stool samples had low diversity with high Proteobacteria and low Bacteroidetes abundance. We demonstrated that, while two milk powders and a legume-based feed were fermentable and produced SCFA in the model, inulin was not fermented to a significant degree. This may reflect the limited microbial diversity, in particular Bacteroidetes needed to ferment longer chain inulin’s. These results demonstrate that assumptions cannot be made regarding the fermentability of carbohydrates by the gut microbiota of children with SAM, and that results obtained in healthy childhood cohorts are not translatable to children with SAM. This is, therefore, an urgent need for future studies screening the fermentability of carbohydrates by the gut microbiota of children with SAM. The results presented in this paper provide insights useful for the development of therapeutic and complimentary feeds for use during treatment and recovery from SAM.

## Materials and Methods

### Substrates

Chicory inulin (catalogue no. I2255) was purchased from Sigma-Aldrich, (Gillingham, UK). Similac Pro-Advance with HMO was purchased from Amazon (UK) and SMA Pro 3 Toddler milk was purchased from Boots Pharmacists (UK). MIMBLE feed was prepared by Campden BRI as described in Walsh et al.(26) Each substrate (0.500 ± 0.005g, dry weight) was weighed into sterilized fermentation bottles (100mL) prior to start of experiment. Nutrient composition (Table S2) was approximated using Nutritics software (Ireland) or using manufacturer provided information.

### Inoculum collection and preparation

Modifying Intestinal Integrity and Microbiome in Malnutrition with Legume-Based Feeds [MIMBLE] was a single centre (Mbale Regional Referral Hospital) randomised comparator trial evaluating safety and feasibility of three feeding strategies (registered on https://pactr.samrc.ac.za as PACTR201805003381361) (26, 27). The protocol was approved by the ethics committees of Imperial College London (15IC3006) and Mbale Regional Referral Hospital (UG-IRC-012). Following parental written consent children were enrolled on day 1 post-admission following consent and followed for 28 days. The trial was conducted to the standards of ICH GCP. Children hospitalised with SAM (n=69) were screened for eligibility (one or more of mid-upper arm circumference (MUAC) <11.5cm, weight-for-height Z-score (WHZ) < -3 or Kwashiorkor) and randomized to either standard milk-feed F75 (n=18); inulin-supplemented standard feeds (InF: n=20) to cowpea supplemented standard feed CPF: n=20). Faecal samples were provided at 7 days following admission to hospital, from trial participants receiving the standard WHO F75/F100 feeding regime and other supportive therapies including antibiotics. Stools from children receiving legume-based feeds were not recruited to this study. Stool samples were frozen and stored at -80°C prior to use. Stool sample was diluted 1:10 with pre-warmed, anaerobic, sterile phosphate buffer saline (0.1M, pH 7.4) in a double meshed stomacher bag (500 mL, Seward, Worthing, UK) and homogenized using a Stomacher 400 (Seward, Worthing, UK) at 200 rpm for 2 cycles at 60 minutes each.

### Batch fermentation

Fermentation experiments were performed with media adapted from Warren and colleagues (2018) (41). In brief, fermentation vessels (100 mL) each contained an aliquot (3.0 mL) of filtered faecal slurry, 82mL of sterilized growth medium, and substrate. The growth medium contained 76mL of basal solution, 5 mL vitamin phosphate and sodium carbonate solution, and 1 mL reducing agent. The composition of the solutions used in the preparation of the growth medium is described in detail in Ravi et al. (42). A single stock (7 litres) of growth medium was used for all vessels prepared for this experiment. Vessel fermentations were pH controlled and maintained at pH 6.8 to 7.2 using 1N NaOH and 1N HCl regulated by a Fermac 260 (Electrolab Biotech, Tewkesbury, UK). A circulating water jacket-maintained vessel temperature at 37°C. A magnetic stirrer was used to keep mixture homogenous and the vessels were continuously sparged with nitrogen (99% purity) maintaining anaerobic conditions. Media with no inoculum was used as a blank whereas, chicory inulin, and SMA, Similac and chickpea supplemented feed with inocula were the experimental conditions evaluated. Samples were collected at 0 (∼5 min post inoculation), 6, 12, 24, 36 and 48 hours after inoculation. The biomass from 1.8 mL aliquot of sample was concentrated by refrigerated centrifugation (4°C; 10,000 g for 10 min), supernatant removed, and then stored at -80°C until DNA extraction. The supernatants were removed to a fresh tube and stored separately at -20°C for ^1^H NMR metabolomic analysis.

### DNA extraction

Concentrated biomass pellets were resuspended in 500 μL (samples collected at 0 and 6 hr) and 650 μL (samples collected at 12 and 24 hr) with sterile, nuclease-free water (Sigma-Aldrich, Gillingham, UK) which was chilled (4°C). The resuspensions were frozen overnight at-80°C, thawed on ice and an aliquot (400uL) was used for bacterial genomic DNA extraction.

FastDNA® Spin Kit for Soil (MP Biomedical, Solon, US) was used according to manufacturer’s instructions which included two bead-beating steps of 60s at a speed of 6.0m/s (FastPrep24, MP Biomedical, Solon, US). DNA concentration was determined using the Quant-iT™ dsDNA Assay Kit, high sensitivity kit (Invitrogen, Loughborough, UK) and quantified on a FLUOstar Optima plate reader (BMG Labtech, Aylesbury, UK).

### Library preparation and 16S rRNA sequencing

Extracted genomic DNA was normalised to 5ng/µl with elution buffer (10mM Tris-HCl). A PCR master mix was made up using 4 ul kapa2G buffer, 0.4 µl dNTP’s, 0.08 µl Polymerase, 0.4 µl 10 µM forward tailed specific primer, 0.4 µl 10 µM reverse tailed specific primer and 12.72 µl PCR grade water (contained in the Kap2G Robust PCR kit Sigma Catalogue No. KK5005) per sample and 18 µl added to each well to be used in a 96-well plate followed by 2 µl of DNA and mixed. Specific PCR conditions were 95LC for 5 minutes, 30 cycles of 95LC for 30s, 55LC for 30s and 72LC for 30 seconds followed by a final 72LC for 5 minutes. Following PCR, a 0.7X SPRI clean-up was performed using KAPA Pure Beads (Roche Catalogue No. 07983298001) eluting the DNA in 20ul of water. A second PCR master mix was made up using 4 ul kapa2G buffer, 0.4 µl dNTP’s, 0.08 µl Polymerase, and 6.52 µl PCR grade water per sample and 11 µl added to each well to be used in a 96-well plate. 2 µl of each P7 and P5 of Nextera XT Index Kit v2 index primers (Illumina Catalogue No. FC-131-2001 to 2004) were added to each well. Finally, the 5 µl of the clean specific PCR was added and mixed. The second PCR was run using 95LC for 5 minutes, 10 cycles of 95LC for 30s, 55LC for 30s and 72LC for 30 seconds followed by a final 72LC for 5 minutes. Final libraries were quantified by Qubit and equimolar pooled together. A single 0.7X SPRI clean-up was performed on the pool. A final Qubit and sizing on High Sensitivity D1000 Screen Tape (Agilent Catalogue No. 5067-5579) using the Agilent Tapestation 4200 was done to calculate the final library pool molarity. The pool was run at a final concentration of 10pM on an Illumina MiSeq instrument using MiSeq® Reagent Kit v3 (600 cycle) (Illumina Catalogue FC-102-3003) following the Illumina recommended denaturation and loading recommendations which included a 20% PhiX spike in (PhiX Control v3 Illumina Catalogue FC-110-3001). The raw data was analysed locally on the MiSeq using MiSeq reporter.

### Bioinformatics analysis

Raw demultiplexed forward and reverse reads were processed using the methods implemented in QIIME2 version 2020.11 with default parameters unless otherwise stated (43). DADA2 was used for paired-end joining, quality filtering, denoising and calling Amplicon Sequence Variants (ASV’s) using the QIIME dada2 denoise-paired method (44). The first 13 bp and the final 50 bp were trimmed before merging due to lower quality scores. The ASV’s were aligned and used to calculate a phylogenetic tree using the QIIME function phylogeny align to tree (45, 46).

Taxonomic assignment of the ASV’s was performed using the QIIME naïve Bayesian classify Scikit-learn using the Silva 99% OTU database (47–49).

The raw sequencing data had a median of 43,909 reads per sample. Following filtering, denoising and merging this was reduced to 37,185 reads per sample. After the removal of chimeric sequences this was further reduced to 32,378 reads per sample. A total of 855 ASV’s were identified across all the samples. Samples with fewer than 1000 reads were excluded from subsequent analysis and the data were rarefied to 17,599 reads. Downstream analysis and visualization was carried out using the PhyloSeq (50) and PhyloSmith (51) packages in the R software package (version 4.0.3), including alpha and beta diversity metrics.

### 1H NMR metabolomics

The samples containing the supernatant from the fermentation media were centrifuged (3,000 x g, 3 min) and 400-μL aliquots were pipetted directly into NMR tubes (Norell® Standard

Series™, 5 mm), followed by the addition of 200 μL of phosphate buffer (NaH_2_PO_4_ (21.7 mM), K_2_HPO_4_ (82.7 mM), NaN_3_ (8.6 mM), 3-(trimethylsilyl)-propionate-d_4_ (TMSP, 1.0 mM), prepared in D_2_O). Spectra were collected on a Bruker NEO 600 MHz spectrometer equipped with a cryoprobe, at a ^1^H frequency of 600 MHz. All experiments were acquired at room temperature, using Bruker’s ‘noesygppr1d’ pulse sequence, with a minimum of 64 scans. A 90° pulse length of 11.09 μs was set for all samples with a mixing time of 0.01 s, acquisition time of 2.62 s, relaxation delay of 4 s, featuring selective pre-saturation (1.0 ms) on the residual H_2_O peak frequency during relaxation delay and mixing time for effective solvent suppression. Spectra were referenced using the TMSP peak (0.0 ppm). The metabolites were quantified using the NMR Suite v7.6 Profiler (Chenomx®, Edmonton, Canada).

### Statistical analysis

Statistical analysis of the metabolite data was carried out in R (version 4.2.0) and SPSS. ANOVA test with post-hoc Tukey’s HSD was used to test for significant differences in metabolite concentrations between substrates and over time.

## Data availability

The raw sequencing data used in this manuscript can be accessed through the NCBI SRA project number PRJNA1080518.

## Acknowledgements

We gratefully acknowledge the technical assistance of David Baker with library preparation and 16S sequencing. We thank all the participants and trial staff participating in the MIMBLE trial.

The authors gratefully acknowledge the support of the Biotechnology and Biological Sciences Research Council (BBSRC); this research was funded by the BBSRC Institute Strategic Programme Food Microbiome and Health BB/X011054/1 and its constituent projects BBS/E/F/000PR13631 and BB/X018857/1.

## References

1. Robertson RC. 2020. The gut microbiome in child malnutrition, p 133–144, Global Landscape of Nutrition Challenges in Infants and Children, vol 93. Karger Publishers.

2. Black RE, Victora CG, Walker SP, Bhutta ZA, Christian P, De Onis M, Ezzati M, Grantham-McGregor S, Katz J, Martorell R. 2013. Maternal and child undernutrition and overweight in low-income and middle-income countries. The lancet 382:427–451.

3. Maitland K, Berkley JA, Shebbe M, Peshu N, English M, Newton CRC. 2006. Children with severe malnutrition: can those at highest risk of death be identified with the WHO protocol? PLoS Med 3:e500.

4. Guideline W. 2013. Updates on the management of severe acute malnutrition in infants and children. Geneva: World Health Organization 2013:6–54.

5. Kerac M, Bunn J, Chagaluka G, Bahwere P, Tomkins A, Collins S, Seal A. 2014. Follow-up of post-discharge growth and mortality after treatment for severe acute malnutrition (FuSAM study): a prospective cohort study. PloS one 9:e96030.

6. Ngari MM, Mwalekwa L, Timbwa M, Hamid F, Ali R, Iversen PO, Fegan GW, Berkley JA. 2018. Changes in susceptibility to life-threatening infections after treatment for complicated severe malnutrition in Kenya. The American journal of clinical nutrition 107:626–634.

7. Qin J, Li R, Raes J, Arumugam M, Burgdorf KS, Manichanh C, Nielsen T, Pons N, Levenez F, Yamada T. 2010. A human gut microbial gene catalogue established by metagenomic sequencing. nature 464:59–65.

8. Walker AW, Duncan SH, Louis P, Flint HJ. 2014. Phylogeny, culturing, and metagenomics of the human gut microbiota. Trends in microbiology 22:267–274.

9. Rowland I, Gibson G, Heinken A, Scott K, Swann J, Thiele I, Tuohy K. 2018. Gut microbiota functions: metabolism of nutrients and other food components. European journal of nutrition 57:1–24.

10. Chambers ES, Preston T, Frost G, Morrison DJ. 2018. Role of gut microbiota-generated short-chain fatty acids in metabolic and cardiovascular health. Current nutrition reports 7:198–206.

11. Spencer SP, Fragiadakis GK, Sonnenburg JL. 2019. Pursuing human-relevant gut microbiota-immune interactions. Immunity 51:225–239.

12. Bhutta ZA, Berkley JA, Bandsma RH, Kerac M, Trehan I, Briend A. 2017. Severe childhood malnutrition. Nature reviews Disease primers 3:1–18.

13. Smith MI, Yatsunenko T, Manary MJ, Trehan I, Mkakosya R, Cheng J, Kau AL, Rich SS, Concannon P, Mychaleckyj JC. 2013. Gut microbiomes of Malawian twin pairs discordant for kwashiorkor. Science 339:548–554.

14. Gordon JI, Dewey KG, Mills DA, Medzhitov RM. 2012. The human gut microbiota and undernutrition. Science translational medicine 4:137ps12–137ps12.

15. Subramanian S, Huq S, Yatsunenko T, Haque R, Mahfuz M, Alam MA, Benezra A, DeStefano J, Meier MF, Muegge BD. 2014. Persistent gut microbiota immaturity in malnourished Bangladeshi children. Nature 510:417–421.

16. Raman AS, Gehrig JL, Venkatesh S, Chang H-W, Hibberd MC, Subramanian S, Kang G, Bessong PO, Lima AA, Kosek MN. 2019. A sparse covarying unit that describes healthy and impaired human gut microbiota development. Science 365:eaau4735.

17. Kristensen KHS, Wiese M, Rytter MJH, Özçam M, Hansen LH, Namusoke H, Friis H, Nielsen DS. 2016. Gut microbiota in children hospitalized with oedematous and non-oedematous severe acute malnutrition in Uganda. PLoS neglected tropical diseases 10:e0004369.

18. Castro-Mejía JL, O’Ferrall S, Krych Ł, O’Mahony E, Namusoke H, Lanyero B, Kot W, Nabukeera-Barungi N, Michaelsen KF, Mølgaard C. 2020. Restitution of gut microbiota in Ugandan children administered with probiotics (Lactobacillus rhamnosus GG and Bifidobacterium animalis subsp. lactis BB-12) during treatment for severe acute malnutrition. Gut Microbes:1–13.

19. Gehrig JL, Venkatesh S, Chang H-W, Hibberd MC, Kung VL, Cheng J, Chen RY, Subramanian S, Cowardin CA, Meier MF. 2019. Effects of microbiota-directed foods in gnotobiotic animals and undernourished children. Science 365:eaau4732.

20. Kerac M, Bunn J, Seal A, Thindwa M, Tomkins A, Sadler K, Bahwere P, Collins S. 2009. Probiotics and prebiotics for severe acute malnutrition (PRONUT study): a double-blind efficacy randomised controlled trial in Malawi. The Lancet 374:136–144.

21. Mostafa I, Nahar NN, Islam MM, Huq S, Mustafa M, Barratt M, Gordon JI, Ahmed T. 2020. Proof-of-concept study of the efficacy of a microbiota-directed complementary food formulation (MDCF) for treating moderate acute malnutrition. BMC Public Health 20:1–7.

22. Johnson CR, Fischer PR. 2020. Feeding the Microbiota: Complementary Foods Enhance Recovery in Malnourished Children by Modulating the Gut Microbiota. Infectious Disease Alert 39.

23. Sheridan PO, Bindels LB, Saulnier DM, Reid G, Nova E, Holmgren K, O’Toole PW, Bunn J, Delzenne N, Scott KP. 2014. Can prebiotics and probiotics improve therapeutic outcomes for undernourished individuals? Taylor & Francis.

24. Logtenberg MJ, Akkerman R, An R, Hermes GD, de Haan BJ, Faas MM, Zoetendal EG, Schols HA, de Vos P. 2020. Fermentation of Chicory Fructo-oligosaccharides and Native Inulin by Infant Faecal Microbiota Attenuates Pro-inflammatory Responses in Immature Dendritic Cells in an Infant-age Dependent and Fructan-Specific Way. Molecular Nutrition & Food Research:2000068.

25. Charbonneau MR, O’Donnell D, Blanton LV, Totten SM, Davis JC, Barratt MJ, Cheng J, Guruge J, Talcott M, Bain JR. 2016. Sialylated milk oligosaccharides promote microbiota-dependent growth in models of infant undernutrition. Cell 164:859–871.

26. Walsh K, Calder N, Olupot-Olupot P, Ssenyondo T, Okiror W, Okalebo CB, Muhindo R, Mpoya A, Holmes E, Marchesi J. 2018. Modifying Intestinal Integrity and Micro Biome in Severe Malnutrition with Legume-Based Feeds (MIMBLE 2.0): protocol for a phase II refined feed and intervention trial. Wellcome open research 3.

27. Calder N, Walsh K, Olupot-Olupot P, Ssenyondo T, Muhindo R, Mpoya A, Brignardello J, Wang X, McKay E, Morrison D. 2021. Modifying gut integrity and microbiome in children with severe acute malnutrition using legume-based feeds (MIMBLE): A pilot trial. Cell Reports Medicine 2:100280.

28. Walsh K, McGurk J, Maitland K, Frost G. 2021. Development of a legume-enriched feed for treatment of severe acute malnutrition. Wellcome Open Research 6.

29. Walsh K, Kiosa A, Olupot-Olupot P, Okiror W, Ssenyond T, Okalebo CB, Muhindo R, Mpoya A, George EC, Frost G. 2023. Modifying gut integrity and microbiome in children with severe acute malnutrition using legume-based feeds (MIMBLE II): A Phase II trial. medRxiv:2023.05. 29.23290673.

30. Koenig JE, Spor A, Scalfone N, Fricker AD, Stombaugh J, Knight R, Angenent LT, Ley RE. 2011. Succession of microbial consortia in the developing infant gut microbiome. Proceedings of the National Academy of Sciences 108:4578–4585.

31. Louis P, Hold GL, Flint HJ. 2014. The gut microbiota, bacterial metabolites and colorectal cancer. Nature reviews microbiology 12:661–672.

32. Caleffi ER, Krausová G, Hyršlová I, Paredes LLR, dos Santos MM, Sassaki GL, Gonçalves RAC, de Oliveira AJB. 2015. Isolation and prebiotic activity of inulin-type fructan extracted from Pfaffia glomerata (Spreng) Pedersen roots. International Journal of Biological Macromolecules 80:392–399.

33. Million M, Raoult D. 2020. Gut dysbiosis in severe acute malnutrition is not an immaturity: The irreversible quantitative-qualitative paradigm shift. Human Microbiome Journal 15:100067.

34. Attia S, Versloot CJ, Voskuijl W, van Vliet SJ, Di Giovanni V, Zhang L, Richardson S, Bourdon C, Netea MG, Berkley JA. 2016. Mortality in children with complicated severe acute malnutrition is related to intestinal and systemic inflammation: an observational cohort study. The American journal of clinical nutrition 104:1441–1449.

35. Hibberd MC, Webber DM, Rodionov DA, Henrissat S, Chen RY, Zhou C, Lynn HM, Wang Y, Chang H-W, Lee EM. 2023. Bioactive glycans in a microbiome-directed food for children with malnutrition. Nature:1–9.

36. Mostafa I, Fahim SM, Das S, Gazi MA, Hasan MM, Saqeeb KN, Mahfuz M, Lynn HB, Barratt MJ, Gordon JI. 2022. Developing shelf-stable Microbiota Directed Complementary Food (MDCF) prototypes for malnourished children: study protocol for a randomized, single-blinded, clinical study. BMC pediatrics 22:1–9.

37. Monira S, Nakamura S, Gotoh K, Izutsu K, Watanabe H, Alam NH, Endtz HP, Cravioto A, Ali S, Nakaya T. 2011. Gut microbiota of healthy and malnourished children in Bangladesh. Frontiers in microbiology 2:228.

38. Kong C, Akkerman R, Klostermann CE, Beukema M, Oerlemans MM, Schols HA, De Vos P. 2021. Distinct fermentation of human milk oligosaccharides 3-FL and LNT2 and GOS/inulin by infant gut microbiota and impact on adhesion of Lactobacillus plantarum WCFS1 to gut epithelial cells. Food & function 12:12513–12525.

39. Falony G, Calmeyn T, Leroy F, De Vuyst L. 2009. Coculture fermentations of Bifidobacterium species and Bacteroides thetaiotaomicron reveal a mechanistic insight into the prebiotic effect of inulin-type fructans. Applied and Environmental Microbiology 75:2312–2319.

40. Blanton LV, Barratt MJ, Charbonneau MR, Ahmed T, Gordon JI. 2016. Childhood undernutrition, the gut microbiota, and microbiota-directed therapeutics. Science 352:1533–1533.

41. Warren FJ, Fukuma NM, Mikkelsen D, Flanagan BM, Williams BA, Lisle AT, Cuív PÓ, Morrison M, Gidley MJ. 2018. Food starch structure impacts gut microbiome composition. Msphere 3.

42. Ravi A, Troncoso-Rey P, Ahn-Jarvis J, Corbin KR, Harris S, Harris H, Aydin A, Kay GL, Le-Viet T, Gilroy R. 2021. Linking carbohydrate structure with function in the human gut microbiome using hybrid metagenome assemblies. bioRxiv.

43. Bolyen E, Rideout JR, Dillon MR, Bokulich NA, Abnet CC, Al-Ghalith GA, Alexander H, Alm EJ, Arumugam M, Asnicar F. 2019. Reproducible, interactive, scalable and extensible microbiome data science using QIIME 2. Nature biotechnology 37:852–857.

44. Callahan BJ, McMurdie PJ, Rosen MJ, Han AW, Johnson AJA, Holmes SP. 2016. DADA2: high-resolution sample inference from Illumina amplicon data. Nature methods 13:581–583.

45. Katoh K, Standley DM. 2013. MAFFT multiple sequence alignment software version 7: improvements in performance and usability. Molecular biology and evolution 30:772–780.

46. Price MN, Dehal PS, Arkin AP. 2009. FastTree: computing large minimum evolution trees with profiles instead of a distance matrix. Molecular biology and evolution 26:1641–1650.

47. Pedregosa F, Varoquaux G, Gramfort A, Michel V, Thirion B, Grisel O, Blondel M, Prettenhofer P, Weiss R, Dubourg V. 2011. Scikit-learn: Machine learning in Python. the Journal of machine Learning research 12:2825–2830.

48. Bokulich NA, Kaehler BD, Rideout JR, Dillon M, Bolyen E, Knight R, Huttley GA, Caporaso JG. 2018. Optimizing taxonomic classification of marker-gene amplicon sequences with QIIME 2’s q2-feature-classifier plugin. Microbiome 6:1–17.

49. Quast C, Pruesse E, Yilmaz P, Gerken J, Schweer T, Yarza P, Peplies J, Glöckner FO. 2012. The SILVA ribosomal RNA gene database project: improved data processing and web-based tools. Nucleic acids research 41:D590–D596.

50. McMurdie PJ, Holmes S. 2013. phyloseq: an R package for reproducible interactive analysis and graphics of microbiome census data. PloS one 8:e61217.

51. Smith S. 2019. Phylosmith: an R-package for reproducible and efficient microbiome analysis with phyloseq-objects. Journal of Open Source Software 4.

