## Supplementary material for "Modelling the gut microbiota of children with malnutrition: *in vitro* models reveal differences in fermentability of widely consumed carbohydrates": Fig. S1

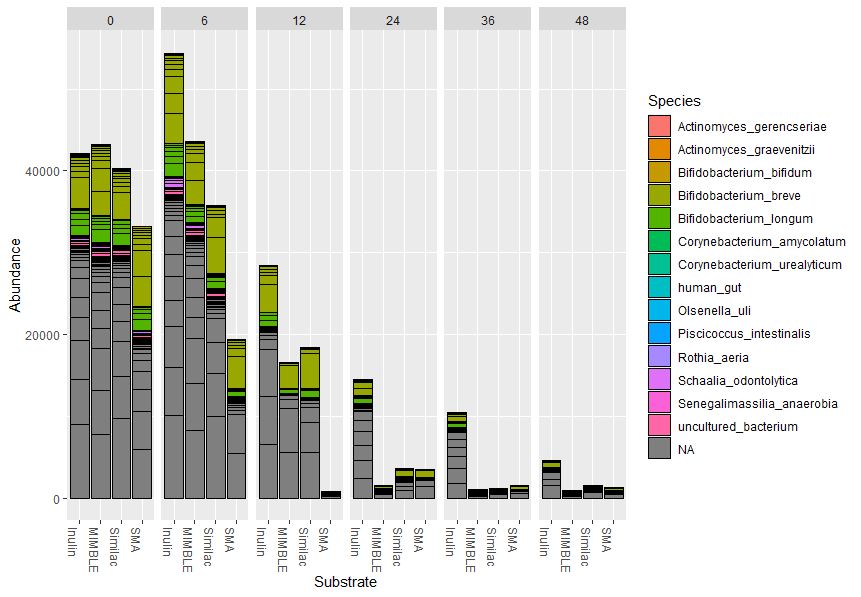

**Figure S1: Abundance of Actinobacteriota in fermentation experiments at 0, 6, 12, 24, 36 and 48 hours of fermentation**

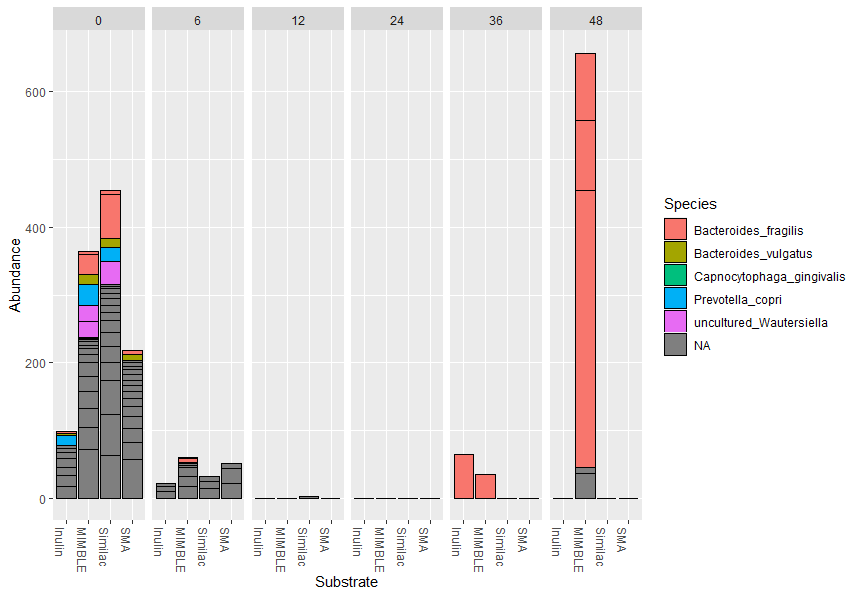

**Figure S2: Abundance of Bacteriodota in fermentation experiments at 0, 6, 12, 24, 36 and 48 hours of fermentation**

**Table S2. Nutritional composition of the substrates used in this study.**

| **Nutrition Information** | **Similac Pro-Advance*** | **SMA Pro 3 Toddler Milk**** | **Mimble**** | **Inulin** |
| --- | --- | --- | --- | --- |
| **Serving Size (mg)** | **500** | **500** | **500** | **500** |
| **Nutrient Composition** |  |  |  |  |
| **Energy (kcal)** | **3.05** | **2.32** | **1.01** | **1.05** |
| **Fat (g)** | **0.17** | **0.12** | **0.06** | **-** |
| **Linoleic Acid (mg)** | **30.49** | **19.09** | **12.50** | **-** |
| **Total Carbohydrates (g)** | **0.32** | **0.25** | **0.09** | **0.49** |
| **Dietary Fiber (g)** | **0.0** | **0.0** | **0.01** | **0.45** |
| **Protein (g)** | **0.06** | **0.05** | **0.02** | **-** |
| **Vitamins**** |  |  |  | **-** |
| **Vitamin A (μg)** | **2.74** | **2.17** | **0.20** | **-** |
| **Vitamin D (μg)** | **0.06** | **0.04** | **0.00** | **-** |
| **Vitamin E (mg)** | **0.03** | **0.03** | **0.01** | **-** |
| **Vitamin K (μg)** | **0.24** | **0.24** | **0.07** | **-** |
| **Thiamin (μg)** | **3.05** | **3.62** | **0.35** | **-** |
| **Riboflavin (μg)** | **4.88** | **9.06** | **0.02** | **-** |
| **Vitamin B6 (μg)** | **1.92** | **2.17** | **0.45** | **-** |
| **Vitamin B12 (μg)** | **0.01** | **0.01** | **0.00** | **-** |
| **Niacin (μg)** | **33.54** | **18.12** | **6.50** | **-** |
| **Folic acid (μg)** | **0.49** | **0.47** | **0.11** | **-** |
| **Pantothenic acid (μg)** | **14.33** | **-** | **1.80** | **-** |
| **Biotin (μg)** | **0.14** | **0.07** | **0.01** | **-** |
| **Vitamin C (mg)** | **0.27** | **0.54** | **0.00** | **-** |
| **Choline (mg)** | **0.73** | **-** | **-** | **-** |
| **Inositol (mg)** | **0.15** | **-** | **-** | **-** |
| **Minerals** |  |  |  | **-** |
| **Calcium (mg)** | **2.50** | **2.90** | **0.50** | **-** |
| **Phosphorus (mg)** | **1.34** | **1.81** | **0.46** | **-** |
| **Magnesium (mg)** | **0.18** | **0.24** | **0.08** | **-** |
| **Iron (mg)** | **0.06** | **0.04** | **0.00** | **-** |
| **Zinc (mg)** | **0.02** | **0.03** | **0.00** | **-** |
| **Manganese (μg)** | **0.15** | **-** | **0.80** | **-** |
| **Copper (μg)** | **2.90** | **1.81** | **0.30** | **-** |
| **Iodine (μg)** | **0.18** | **0.43** | **0.05** | **-** |
| **Selenium (μg)** | **0.06** | **0.05** | **0.01** | **-** |
| **Sodium (mg)** | **0.76** | **1.01** | **0.20** | **-** |
| **Potassium (mg)** | **3.35** | **3.26** | **0.73** | **-** |
| **Chloride (mg)** | **2.07** | **1.52** | **0.42** | **-** |
| ***Nutrient from manufacturer**  ****Nutrient calculated from Nutritics software** | |  |  |  |
